# Co-limitation by stable, dynamic and directional habitat features shapes climate vulnerability in an alpine specialist

**DOI:** 10.64898/2026.04.09.717579

**Authors:** Timothy M. Brown, Reza Goljani Amirkhiz, Sarah Albright, Alena Arnold, Emma Brown, Carl Brown, Vincent Chevreuil, Ryan Cheung, Daisy Cortes, Jessica Gallardo, Karim Hanna, Raquel Rodriguez Lozano, Juan Rebellon, Leticia Santillana, Kevin Silberberg, Juyung Yoo, Kathryn Bernier, Kristen C. Ruegg, Mevin B. Hooten, Erika S. Zavaleta

**Affiliations:** Ecology and Evolutionary Biology Department, University of California, Santa Cruz, CA, USA; Department of Biology, Colorado State University, Fort Collins, CO, USA; Department of Statistics and Data Sciences, The University of Texas at Austin, Austin, TX, USA

**Keywords:** alpine birds, habitat selection, climate change, snowpack, woody encroachment, species distribution, abundance, co-limitation, Sierra Nevada, *Leucosticte tephrocotis dawsoni*

## Abstract

Alpine ecosystems are among the most climate sensitive on Earth, yet logistical challenges and detection biases often impede robust assessment of alpine dependent species. We investigated habitat associations and density patterns of the Sierra Nevada subspecies of the Gray-crowned Rosy-Finch (*Leucosticte tephrocotis dawsoni*), an alpine obligate and regional endemic, over five breeding seasons from 2018 to 2022 using hierarchical distance sampling and mark–recapture distance sampling to explicitly account for imperfect detection and spatial heterogeneity. Density estimates tracked annual snowpack variation, ranging from 4.77 individuals/km² in a low snow year to 12.08 individuals/km² in a high snow year. Abundance was highest near persistent snow patches that provide foraging habitat and near cliffs that provide nesting substrate, and declined sharply above approximately 10% woody cover, with densities approaching zero beyond approximately 25%, indicating a steep ecological threshold. In contrast, the proportion of surveyed blocks with detections remained relatively stable across years. Together, these patterns indicate a three timescale co-limitation framework in which breeding habitat is shaped by static features (cliffs), dynamic annual drivers (snowpack), and longer-term directional change (woody encroachment). By linking population density to climate sensitive habitat features, this study provides a high-resolution abundance-based baseline for long term monitoring and offers a framework for evaluating climate vulnerability in alpine and other resource-limited systems.

**Open Research Statement:** Data necessary to replicate the analyses and results presented in this manuscript will be archived in the Dryad Digital Repository upon acceptance, with no embargo on the material.

The R code associated with this manuscript is not novel. All analyses use publicly available packages and functions without modification, including mrds (v2.3.0), unmarked (v1.2.5), tidyverse (v2.0.0), MuMIn (v1.47.5), and standard ggplot2 visualization tools. All code is properly cited within the manuscript and publicly available through CRAN. Complete analysis scripts will be archived in Dryad alongside the data upon acceptance to facilitate full reproducibility of results.

## Introduction

Alpine species are among the most climate-sensitive organisms on Earth, making them critical early indicators of ecological change (Gottfried et al. 2012; Pepin et al. 2015). Rapid, directional climate change underscores the urgency of establishing ecological baselines and tracking species abundance over time, particularly for range-restricted alpine taxa whose dynamics can act as sentinels for broader ecosystem change (Moritz et al. 2008; Tingley et al. 2009; Beever et al. 2012; Pauli et al. 2012). Compared to occupancy alone, abundance data offer earlier and more sensitive indicators of ecological change, often detecting stress responses before extirpation occurs (Joseph et al. 2006; Magurran et al. 2010; Beever et al. 2012; Lehikoinen et al. 2019; Rumpf et al. 2019). This makes abundance essential for conservation prioritization and adaptive management under climate change (Nielsen et al. 2009; Beever et al. 2013). Understanding spatial and temporal variation in abundance is foundational to ecological research and provides critical insight for conservation and management in the face of environmental shifts (Buckland et al. 2004; Nielsen et al. 2005; Pauli et al. 2012; Keith et al. 2013; Pacifici et al. 2017; Bowler et al. 2019). However, many species inhabit remote or challenging environments where accurate abundance measurements are hindered by detection biases and observation error (Yoccoz et al. 2001; Andersen & Steidl 2020).

Range-restricted alpine species, often understudied due to remoteness, offer critical insights into climate sensitivity (Socolar et al. 2017). Alpine ecosystems are warming rapidly, with altered snowpack dynamics driving biological change (Grabherr et al. 2010; Pauli et al. 2012; Pepin et al. 2015; Siirila-Woodburn et al. 2021). These systems are among the most climate-vulnerable on Earth, with broad taxonomic impacts now well documented (Parmesan & Yohe 2003; Moritz et al. 2008; Gottfried et al. 2012; Rumpf et al. 2019). Alpine specialists, in particular, are sensitive to variations in temperature, precipitation, and snowpack dynamics, as these directly influence habitat quality, prey availability, and reproductive success (Erb et al. 2011; Mason et al. 2014; García-González et al. 2016; Resano-Mayor et al. 2019; De Gabriel Hernando et al. 2022; Green et al. 2022; Winkler et al. 2023). Climate-driven alterations in these factors significantly reduce habitat suitability, exacerbate fragmentation, and potentially disrupt population connectivity and persistence (Schoville et al. 2018; Scridel et al. 2018; Rumpf et al. 2019; Brambilla et al. 2022b; De Gabriel Hernando et al. 2022). For example, recent studies of European alpine birds demonstrate how declining snowpack and rising temperatures reduce habitat connectivity and increase local extirpation risks (Lehikoinen et al. 2019; Resano-Mayor et al. 2019; Brambilla et al. 2022a). These ecosystems pose significant research challenges due to rugged terrain, low productivity, and patchy species distributions, yet they remain critical barometers of climate vulnerability (Pepin et al. 2015; Engler et al. 2017; García-Navas et al. 2020).

Recent studies also emphasized the need for rigorous abundance data to quantify habitat associations accurately, especially for range-restricted alpine species facing rapid climate-driven habitat alteration (Beever et al. 2012; Socolar et al. 2017; De Gabriel Hernando et al. 2022). To elucidate how factors shaping abundance at fine spatial scales and across years inform understanding of current and future climate vulnerability of alpine species, we sought to characterize breeding habitat requirements of one of North America’s highest breeding birds, the Sierra Nevada Gray-crowned Rosy-Finch (*Leucosticte tephrocotis dawsoni*; hereafter, SN Rosy-Finch). The SN Rosy-Finch is an ideal indicator for evaluating alpine climate vulnerability due to its limited elevational range, specialized reliance on climate-sensitive resources such as persistent snowfields, and the high detectability afforded by its open, sparsely vegetated habitat. Despite being a key indicator species for alpine ecosystem health, this taxon remains one of North America’s least-studied passerines, with sparse data regarding its specific habitat requirements and climate vulnerability (Grinnell & Storer 1924; Twining 1940; Stanek 2009; Macdougall-Shackleton et al. 2020), though see Epanchin et al. (2010) for a notable exception focused on foraging ecology, and Tingley et al. (2009) for broader evidence of climate-driven range shifts in the Sierra Nevada avifauna. Modeling of historical habitat suitability indicates substantial range contraction and fragmentation for SN Rosy-Finch, consistent with broader trends observed in alpine birds experiencing elevational range contractions globally (Freeman et al. 2018; Scridel et al. 2018; Golijani Amirkhiz et al. 2025). This underscores the need for contemporary data to inform conservation strategies. For the SN Rosy-Finch, the southernmost taxon of the North American rosy-finch species complex, previous work has shown that cliffs and snow patches are critical nesting and foraging resources (Epanchin et al. 2010). Similar patterns have been documented in other high-alpine passerines, where snowfields provide seasonal pulses of arthropod prey that shape breeding-season habitat use (Antor 1995; Scridel et al. 2018; Brambilla et al. 2019). Despite this, little is known about how spatial and temporal variation in such fixed (e.g., cliffs as potential nest sites) and dynamic (e.g., summer snowpack, woody encroachment) habitat features influence abundance across the subspecies’ range. We addressed this gap through a five-year study spanning dramatic interannual variation in climate and summer snowpack, explicitly evaluating the relative roles of these features in shaping SN Rosy-Finch abundance patterns.

Summer snowpack is a key ecological resource in alpine ecosystems because melting snowfields concentrate invertebrate prey along their edges, forming ephemeral but highly productive foraging zones during the brief alpine breeding season (Antor 1995; Epanchin et al. 2010). However, declining snowpack trends and earlier snowmelt periods (Mote et al. 2018; Siirila-Woodburn et al. 2021) could reduce the reliability of these foraging resources, necessitating greater foraging flexibility and potentially larger home ranges. Recent projections specific to the Sierra Nevada indicate that April 1 snow water equivalent could decline by approximately 76 percent under a plus-three degree Celsius warming scenario, with the region transitioning from a snow-dominated to a rain-dominated hydrological regime (Beltrán-Peña et al. 2025). Fine-scale variability in snowpack extent and duration alters the timing and spatial availability of snowmelt-foraging zones and has been shown to drive within-season shifts in alpine bird distributions, highlighting the need for spatially and temporally explicit habitat modeling (Ceresa et al. 2020; Brambilla et al. 2022b). Studies of other alpine-breeding birds indicate that declining snowpack reduces these foraging opportunities by eliminating the moist, cool microclimates that sustain insect emergence, ultimately limiting chick provisioning and reducing reproductive success (Resano-Mayor et al. 2019; Brambilla et al. 2022a).

In contrast to Rocky Mountain alpine ecosystems, where higher and more consistent summer rainfall supports stable, productive habitats (Bueno de Mesquita et al. 2018), California’s alpine habitats experience patchier and more ephemeral snowfields due to their drier Mediterranean climate (Siirila-Woodburn et al. 2021). These differences suggest that SN Rosy-Finch may experience greater resource unpredictability and scarcity compared to closely related Rocky Mountain and Intermountain West taxa, as indicated by studies of Brown-capped Rosy-Finches in Colorado (Bernier et al. 2023, *Colorado Parks and Wildlife Technical Report*), Black Rosy-Finches in the Intermountain West (Brown et al. 2018), and prior work on SN Rosy-Finches in California (Epanchin et al. 2010), potentially influencing their vulnerability to climate change.

In addition to interannual snowpack variability, gradual encroachment of woody vegetation driven by warming temperatures and declining snowpack may further constrain historically open breeding habitats for alpine specialists (Lubetkin et al. 2017; Scridel et al. 2018). Some alpine bird species show negative responses to increased shrub and conifer cover, which can reduce habitat openness, limit visibility, and increase predation risk (Braunisch et al. 2016; Chamberlain et al. 2016; Scridel et al. 2018). Globally, conifer encroachment into alpine ecosystems is accelerating (Dolanc et al. 2013; Macias-Fauria and Johnson 2013), with documented shifts in subalpine forest structure and upward expansion in the Sierra Nevada over the past century, raising concern that SN Rosy-Finches may be vulnerable to ongoing habitat change. To evaluate these possibilities, we asked: To what extent do cliffs, persistent snowfields, and woody vegetation structure abundance patterns of SN Rosy-Finch across their elevational and geographic range in California?

To test our hypotheses that cliffs, snowpack, and woody vegetation co-limit breeding habitat availability for SN Rosy-Finches, and that variation in snowpack and woody encroachment trends may increase their vulnerability to climate change, we used hierarchical distance sampling methods over five consecutive breeding seasons from 2018 to 2022, capturing interannual variability in snowpack and habitat conditions across the Sierra Nevada, White Mountains, and Sweetwater Mountains in California, USA. This approach explicitly addresses detection biases and spatial heterogeneity, ensuring robust density estimates critical for conservation management (Buckland et al. 2001; Royle et al. 2004a). We hypothesized that Rosy-Finch abundance is co-limited by fixed (cliffs), dynamic (snowpack), and directional (woody vegetation) habitat components, each responding to climate change at distinct temporal scales.

## METHODS

### Study Area

#### Alpine Ecosystem Characteristics

Our study was conducted across alpine ecosystems in the Sierra Nevada, White Mountains, and Sweetwater Mountains in California, USA. Alpine zones are defined as habitats above the treeline, characterized by harsh climatic conditions, short growing seasons, and specialized ecological communities (Körner & Paulsen 2004; Rundel 2011). In California, these alpine ecosystems experience short winter precipitation periods followed by prolonged summer droughts typical of the Mediterranean climate, limiting snow persistence and water availability in the summer months (Rundel & Millar 2016; Siirila-Woodburn et al. 2021). Recent studies suggest these Mediterranean alpine ecosystems are particularly susceptible to increasing climatic extremes, making them ideal systems for examining ecological responses to climate variability (Pepin et al. 2015; Mote et al. 2018).

#### Focal Species Ecology

The SN Rosy-Finch is endemic to alpine environments in California and western Nevada, where the Mediterranean climate limits snow retention and reduces summer rainfall (Mote et al. 2018; Siirila-Woodburn et al. 2021). In these exposed, high-elevation landscapes, the species relies on isolated microhabitats such as rocky cliffs (nesting), snowfields (foraging), and alpine vegetation patches (foraging and predator avoidance) (Stanek 2009; Brown et al. 2018; Bernier et al. 2023). Rosy-Finches forage on a broad mix of alpine resources, including seeds, flowers, and leaves from tundra plants, as well as insects and other invertebrates that accumulate on snow surfaces or emerge from aquatic and wetland areas (Macdougall-Shackleton et al. 2000; Epanchin et al. 2010). This ecological flexibility allows them to exploit patchy and ephemeral food sources, although their overall range remains highly restricted to cold, open, high-elevation terrain.

#### Regional Variation Among Mountain Ranges

The Sierra Nevada extends approximately 600 km north-to-south, reaching its maximum elevation at Mount Whitney (4,421 m), the highest peak in the contiguous United States. The Sierra Nevada supports extensive alpine habitats shaped historically by glacial processes, characterized by comparatively higher annual precipitation and more persistent snowpack relative to adjacent ranges (Millar et al. 2004; Rundel & Millar 2016). In contrast, the White Mountains (4,344 m max elevation) and Sweetwater Mountains (3,552 m max elevation) lie east of the Sierra Nevada, closer to the Great Basin Desert. These ranges experience more pronounced aridity, lower productivity, reduced snow persistence, and greater climatic variability driven by convective summer storms rather than stable snow accumulation, resulting in patchier resource distributions and distinct ecological communities (Beever & Brussard 2004; Millar et al. 2018). Historical analyses demonstrate that climatic gradients structure regional SN Rosy-Finch distributions across mountain ranges (Goljani Amirkhiz et al. 2025), with Sierra Nevada populations occupying distinct thermal and precipitation niches relative to populations in the White and Sweetwater Mountains. These climatic differences reinforce the importance of examining habitat associations across mountain ranges to understand region-specific climate vulnerability.

### Site selection and survey design

We identified potential sampling sites using a spatially stratified random design to capture variation across the species’ elevational and geographic range. Site selection was based on elevation (>2,500 m) and terrain ruggedness quantified using the Vector Ruggedness Measure (VRM; Sappington et al. 2007), a unitless index of three-dimensional surface complexity derived from a 30 m digital elevation model. Cells with VRM ≥ 0.03 were classified as cliff or rugged terrain in a binary habitat layer, a threshold validated against field-confirmed cliff locations across multiple sites and years. Elevation and VRM layers were intersected in ArcGIS Pro 3.3.1 (Esri, 2023) to produce a binary “suitable habitat” raster representing areas with high likelihood of supporting SN Rosy-Finch nesting and foraging. The proportion of suitable habitat within each candidate 4 × 4 km sampling block was calculated from this raster and used to stratify blocks as described below. Slope was evaluated as a continuous terrain attribute during preliminary GIS screening and later included as a covariate in detection models, but was not used as a threshold-based filter in defining suitable habitat. To incorporate emerging insights on alpine bird occupancy, we also evaluated landscape-scale context, including patch size, terrain heterogeneity, and proximity to persistent snowfields (Brambilla et al. 2017; De Gabriel Hernando et al. 2022).

Sampling blocks measured 4 × 4 km and were stratified by the proportion of suitable habitat, prioritizing blocks with ≥75% suitable habitat within 6 km of trail or road access to reflect logistical constraints of alpine fieldwork (Ralph et al. 1993; Buckland et al. 2001). We also included blocks with moderate (50–74%) and low (25–49%) habitat coverage to ensure representation across a full suitability gradient. This process yielded 27 sampling blocks spanning the Sierra Nevada, White Mountains, and Sweetwater Mountains (Fig. 1a). Within each block, we established 2-km transects placed randomly within 45° of an origin point at the block center or edge, maintaining a ≥125 m buffer from block boundaries to avoid edge effects (Fig. 1b). Fixed-point survey stations were positioned every 250 m along transects to maintain spatial independence and minimize double counts, consistent with distance sampling best practices (Fig. 1b, Fig. S1; Buckland et al. 2001; Thomas et al. 2010). The 250 m spacing exceeded twice the effective detection radius estimated from the MRDS analysis (EDR = 94.1 m), ensuring that detection zones of adjacent stations did not overlap and minimizing the potential for double-counting individuals between consecutive points.

**Figure 1.**
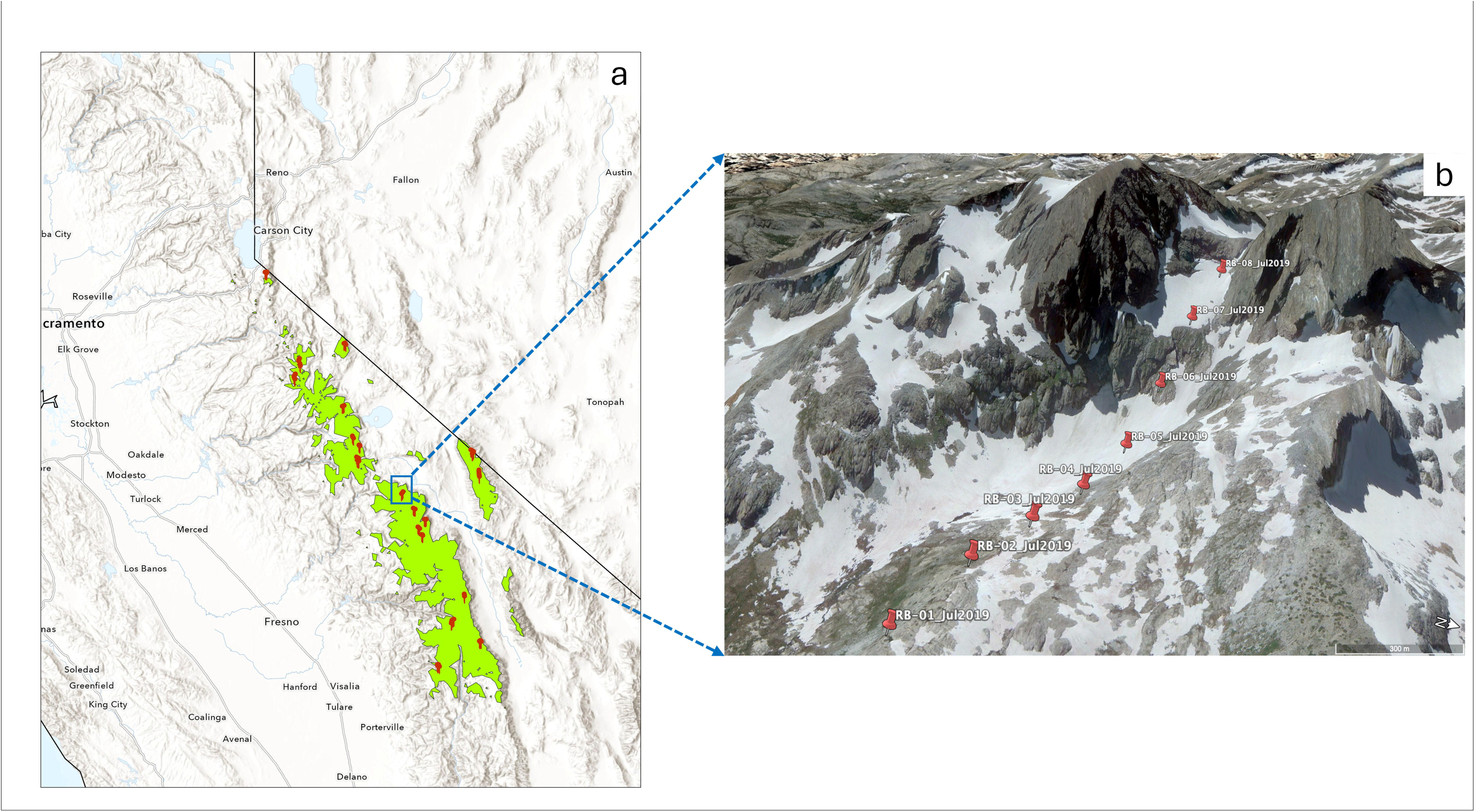
Study area and survey design. (a) Predicted suitable breeding habitat for Sierra Nevada Rosy-Finch (green shading; elevations ≥2,900 m) across California, with 4×4-km sampling blocks (red symbols) spanning the Sierra Nevada, White Mountains, and Sweetwater Mountains. (b) Aerial view of a Sierra Nevada sampling block at Mt. Ritter–Banner Peak (July 2019), showing point-count stations RB-01 through RB-08 (red markers) spaced every 250 m along a survey transect. The image illustrates co-occurring cliff faces (nesting substrate) and persistent snowfields (foraging habitat) characteristic of surveyed alpine terrain. Scale bar: 300 m. See Methods for survey design details and Table S1 for annual survey effort by block.

### Bird surveys

We conducted point-count surveys using a double-observer, independent distance sampling protocol (Nichols et al. 2000; Buckland et al. 2001; Fletcher and Hutto 2006) during the breeding season from late May to early August across five years (2018–2022). Surveys were completed between one hour after sunrise and midday (0600–1300 h), avoiding later periods of reduced song activity. Across the study, we surveyed a total of 216 point-count stations, with each visited three to five times per breeding season; coverage expanded from 128 stations in 2018 to 168 by 2022. Annual variation in snowpack and precipitation provided a natural experimental gradient, ranging from record-high to record-low conditions (38–162% of long-term averages; California DWR 2023), allowing robust inference on habitat associations under contrasting climatic scenarios (Erb et al. 2011; García-González et al. 2016; Resano-Mayor et al. 2019).

To estimate detection probability and correct for imperfect detection, we used a mark–recapture distance sampling (MRDS) framework (Laake & Borchers 2004), which integrates distance sampling with a double-observer design to estimate detection at distance zero (g(0)). At each station (see Fig. 1b), two observers independently conducted 7-min counts, recording all SN Rosy-Finches detected visually or aurally, and measuring radial distances with a laser rangefinder (Leupold RX-1600i). Observations beyond 165 m were excluded based on detection histograms showing sharply reduced detection probability at greater distances, consistent with best practices in avian distance sampling (Buckland et al. 2001; Thomas et al. 2010).

Juveniles/fledglings were excluded to avoid age-related detection biases. To reduce observer effects, we employed an independent double-observer protocol with temporally staggered arrivals (3–5 min apart) due to steep alpine terrain, ensuring no visual or verbal interaction and maintaining separate field data sheets. This staggered approach is supported in challenging landscapes (Fletcher and Hutto 2006) and preserved the MRDS assumption of observer independence. Estimated observer-specific detection probabilities were high (primary = 0.88, secondary = 0.86; Table 1), consistent with ecological expectations for a low-density, centrally foraging, and site-faithful alpine specialist, lending confidence to our estimates. Observer-specific detection probabilities and full MRDS model results are presented in the Results section (Table 1).

**TABLE 1.** Detection metrics from independent double-observer mark–recapture distance sampling for Sierra Nevada Rosy-Finch (2018–2022).

| Metric | Value | Interpretation |
| --- | --- | --- |
| Single Observer $p(0)$ | 0.87 | Probability of detecting a bird by one observer |
| Combined $p(0)$ | 0.98 | Detection probability when both observers are combined |
| EDR (m) | 94.10 | Effective distance within which detection is reliable |
| Density (birds/km<sup>2</sup>) | 72.60 | Average density in occupied areas (where detections occurred) |
| Survey Area (ha) | 573.05 | Total area covered in MRDS analysis (165 m truncation) |

*Note:* Models fit using the mrds R package with a 165 m truncation distance. Abbreviations: $p_0$ , observer-specific detection probability; $p()$ , detection probability at distance zero; EDR, effective detection radius (m).

We measured distances from each survey point to the nearest cliff, persistent snow patch, and water body to characterize key habitat features for nesting and foraging. Cliffs were defined as slopes ≥30°, snow patches as snowfields that remained present through at least one full survey visit within the breeding season (late May–early August) and were of sufficient extent to support active foraging observations, and water bodies as lakes, ponds, or wetlands. These features have been identified as critical predictors of alpine passerine breeding habitat in other mountain ecosystems (Brambilla et al. 2017; Resano-Mayor et al. 2019).

### Habitat surveys

Vegetation and habitat surveys were conducted from 2019 to 2022 within 50-m radius plots centered at each point-count station, following a modified relevé protocol specifically adapted for alpine environments, where patchy vegetation, steep slopes, and late-lying snow can obscure microhabitat boundaries (Ralph et al. 1993). The 50-m radius was selected to match the spatial scale at which SN Rosy-Finch nesting and foraging habitat features can be reliably detected while maintaining comparability with other alpine bird studies.

Each plot was classified into ten distinct habitat types: mixed tundra, meadow tundra, prostrate shrub tundra, shrub-dominated wetland/riparian, graminoid/forb-dominated wetland, woody vegetation (conifers), bare ground tundra, exposed rock, snow, and water. Water bodies were further categorized by flow and permanence (lotic, ponds, lakes) to capture differences relevant to foraging and thermoregulation opportunities. Human disturbances (trails, campsites, roads, mining) were documented because such features can alter alpine bird behavior and habitat use.

Vegetation cover was visually estimated using structured cover categories (1%, 2%, 5%, 10–90%), a method shown to reduce observer bias and increase repeatability in alpine vegetation assessments (Pavlacky et al. 2017). All observers received pre-season calibration training to ensure consistency across years and surveyors. This classification scheme allowed for direct integration of vegetation metrics with distance-to-feature variables (cliffs, snow, water) in subsequent analyses, ensuring that both structural and compositional habitat attributes were captured at scales biologically relevant to the focal species.

### Analytical framework

We estimated SN Rosy-Finch abundance a three-step analytical approach consisting of Generalized Linear Mixed Models (GLMM), Mark–Recapture Distance Sampling (MRDS), and Hierarchical Distance Sampling (HDS). These approaches were applied sequentially. GLMMs were used to evaluate whether spatial (block) or temporal (year) grouping better explained heterogeneity in raw counts; MRDS was used to estimate detection probability at distance zero and evaluate observer effects in a dual-observer framework; and HDS was used to estimate detection-corrected abundance as a function of habitat covariates while modeling detection through distance intervals. To evaluate whether occupancy-based metrics would have detected the same interannual variation as abundance estimates, we compared annual detection frequencies (proportion of surveyed blocks with ≥1 detection; Table S1) against year-specific density estimates from HDS models (Table 2). This comparison was post hoc because a more formal inference would require a hierarchical occupancy model.

**TABLE 2.** Annual density estimates for Sierra Nevada Rosy-Finch from hierarchical distance sampling models, 2018–2022.

| Year | $\lambda$ (ind/km<sup>2</sup>) | SE | 95% CI (lower) | 95% CI (upper) |
| --- | --- | --- | --- | --- |
| 2018 | 4.77 | 1.69 | 2.12 | 8.31 |
| 2018 | 4.95 | 1.96 | 1.85 | 9.78 |
| 2018 | 4.69 | 1.30 | 2.60 | 7.35 |
| 2019 | 12.08 | 2.68 | 6.96 | 17.63 |
| 2020 | 7.35 | 1.70 | 4.55 | 10.84 |
| 2020 | 7.58 | 2.16 | 3.90 | 12.19 |
| 2021 | 7.75 | 2.18 | 3.37 | 11.88 |
| 2021 | 7.82 | 2.33 | 4.17 | 12.42 |
| 2021 | 7.42 | 2.22 | 3.72 | 12.90 |
| 2022 | 5.01 | 1.25 | 2.81 | 7.78 |

*Note:* Densities estimated from HDS fitted separately by year using a negative binomial distribution; uncertainty reflects 200-iteration parametric bootstrap 95% confidence intervals. Values expressed per km<sup>2</sup> of surveyed alpine habitat. Annual estimates from best-supported model (lowest AIC); supported model set ( $\Delta\text{AIC} \leq 2$ ) reported in Table 3. Survey effort summarized in Table S1. Abbreviations: $\lambda$ , density (individuals per km<sup>2</sup>); SE, standard error; CI, confidence interval; HDS, hierarchical distance sampling.

#### GLMM Analysis

Prior to hierarchical distance sampling, we used GLMMs (glmmTMB package; Brooks et al. 2017) to evaluate whether spatial (block) or temporal (year) grouping structure better explained heterogeneity in raw count data. Because *unmarked* does not support simultaneous random effects for block and year, this comparison informed the random-effect structure used in subsequent HDS models. We compared models with identical fixed effects (selected *a priori* based on Rosy-Finch ecology) but differing random-effect structure: (1|Block), (1|Year), or no random effect. Counts were modeled with a negative binomial distribution to address overdispersion (Lindén and Mäntyniemi 2011), and covariates were Z-score standardized (mean = 0, SD = 1).

#### MRDS Analysis

To evaluate observer-specific detectability and estimate detection probability at distance zero, we fit MRDS models (mrds R package; Laake et al. 2017) using a 165 m truncation to reduce the influence of sparse detections at larger distances (Buckland et al. 2001; Thomas et al. 2010). This cutoff corresponded to approximately the 95th percentile of observations and follows common truncation guidance for avian distance sampling (Buckland et al. 2015). Although mark–recapture is commonly associated with individually marked animals,

MRDS can be applied to unmarked populations in dual-observer surveys by treating detections by one observer as “marks” for the second observer (Nichols et al. 2000; Buckland et al. 2001, 2004). We used the independent observer (io) method (Laake & Borchers 2004), which assumes observers operate independently, conditional on distance.

We selected a half-normal key function *a priori* because we expected a smooth monotonic decline in detectability with distance for point counts in open alpine terrain, and we compared it to a hazard-rate key function. We also explored detection models, including a limited set of additional covariates (Julian day, time of day, slope); these did not improve AIC relative to the distance-only half-normal model. The top-ranked model indicated that distance alone adequately described detection (p(0) = 0.98, 95% CI: 0.96–0.99; effective detection radius = 94.1 m), with similar detectability between observers (primary = 0.88; secondary = 0.86). Because observer identity did not improve model fit, detections from both observers were pooled before HDS analyses. HDS models detection through distance intervals within the *gdistsamp* likelihood rather than observer-specific covariates; therefore, pooling observers was appropriate after observer effects were shown to be minimal. Full MRDS protocols, model comparisons, and diagnostics are provided in Supplementary Materials (Table S2, Fig. S2).

#### HDS Analysis

We estimated detection-corrected abundance using hierarchical distance sampling implemented via the gdistsamp() function in the unmarked R package (Fiske & Chandler 2011). HDS jointly estimates detection probability (scale parameter σ) and abundance (expected count λ), linking abundance to site-level habitat covariates while modeling detection through discrete distance intervals in the likelihood rather than including continuous distance as a covariate. We specified a negative binomial mixture to account for overdispersion (Lindén and Mäntyniemi 2011). Abundance covariates included distance to cliffs, snow cover, woody vegetation, and a snow × cliff interaction, whereas detection covariates included Julian day, slope, and time of day. The snow × cliff interaction tested the hypothesis that breeding density is highest where nesting substrate (cliffs) and snow-associated foraging resources co-occur, consistent with potential joint resource limitation. All covariates were Z-score standardized (mean = 0, SD = 1).

Models were fit separately for each year (2018–2022) for three reasons: preliminary GLMMs indicated stronger spatial heterogeneity among blocks than temporal variation among years (see Results), *unmarked* does not support simultaneous block and year random effects, and our objective was to evaluate whether density–habitat relationships shifted across contrasting snowpack years, which is most directly assessed using year-specific models. For each year, we evaluated 3–8 candidate models representing a priori ecological hypotheses rather than all possible covariate combinations. Candidate sets included (i) core resource models (distance to snow, distance to cliffs, woody vegetation), (ii) interaction models including snow × cliffs, (iii) models adding slope or distance to water, and (iv) reduced single-resource models. Model selection was based on AIC, with ΔAIC ≤ 2 indicating equivalent support (Burnham & Anderson 2002). Parameter uncertainty was quantified using 200-iteration parametric bootstrapping to generate standard errors and 95% confidence intervals.

## Results

### Model Validation and Performance

Mark-Recapture Distance Sampling (MRDS) models indicated consistently high detection probabilities for SN Rosy-Finch, confirming that survey methods and protocols were well-suited for monitoring this high-elevation, sparsely distributed species. The combined detection probability at zero distance was very high (p(0) = 0.98, 95% CI: 0.96–0.99; Table 1), indicating robust detectability. This was further supported by a high Effective Detection Radius (EDR = 94.1 m), signifying that detections remained reliably high within this distance threshold (Fig. S1). Goodness-of-fit assessments confirmed model robustness, with chi-square tests (χ² = 20.82, p = 0.79) and Freeman-Tukey residual analyses indicating strong model performance (Table S2). We observed slight overdispersion (ĉ = 1.25), explicitly indicating modest unmodeled heterogeneity. However, this value was within the acceptable range (ĉ < 2) typically reported for ecological count data analyzed by hierarchical modeling (Kéry & Royle 2016), confirming that our models accounted for key sources of variability. This modest overdispersion is consistent with clustered bird distributions in alpine environments (e.g., Brambilla et al. 2017; Resano-Mayor et al. 2019) and does not indicate systematic model misspecification (Kéry & Royle 2016). Parametric bootstrapping further validated MRDS results, with observed fit statistics consistently falling within their respective 95% confidence intervals (Table S2). Observer detection probabilities were consistently high (primary = 0.88; secondary = 0.86; Table 1), indicating strong agreement and minimal observer-related bias, providing confidence in the detection process and ensuring reliable inputs for subsequent HDS-based abundance estimates.

### Random-Effect Structure

Preliminary GLMM comparisons indicated that models including a block-level random effect substantially outperformed models including a year-level random effect (AIC = 853.3 vs. 867.6; ΔAIC = 14.3) and fixed-effects-only models (ΔAIC = 10.3), indicating stronger spatial heterogeneity among survey blocks than temporal variation among years. These results supported retaining block as the grouping structure in abundance modeling while fitting hierarchical distance sampling models separately by year to evaluate interannual variation in density–habitat relationships.

Hierarchical Distance Sampling (HDS) models consistently preferred negative binomial over Poisson distributions, reflecting common overdispersion in ecological count data due to environmental heterogeneity and clustered bird distributions (Burnham & Anderson 2002). Annual ĉ values ranged from 1.01 to 1.94, with the highest in 2019 (Table S2). Detection probabilities increased with Julian day (Table S3) and declined with increasing slope steepness (Table S3). Detection probability followed a quadratic relationship with time of day, peaking during mid-morning surveys (approximately 09:00–10:30) and declining through the afternoon, consistent with typical alpine passerine activity patterns (Fig. S3). These patterns align with ecological expectations for alpine passerines (Epanchin et al. 2010; Brambilla et al. 2017).

### Density Estimates and Ecological Covariates

Annual HDS density estimates for SN Rosy-Finch varied across the study period, with lowest values in 2018 (4.77 individuals/km²; 52% of historical snowpack) and highest in 2019 (12.08 individuals/km²; 162% snowpack), and intermediate densities in 2020–2022 (Table 2, Fig. 3). Standard errors ranged from 1.45 to 2.62 individuals/km², and 95% confidence intervals were narrow relative to effect sizes, indicating stable estimation precision (Table 2).

**Figure 2.**
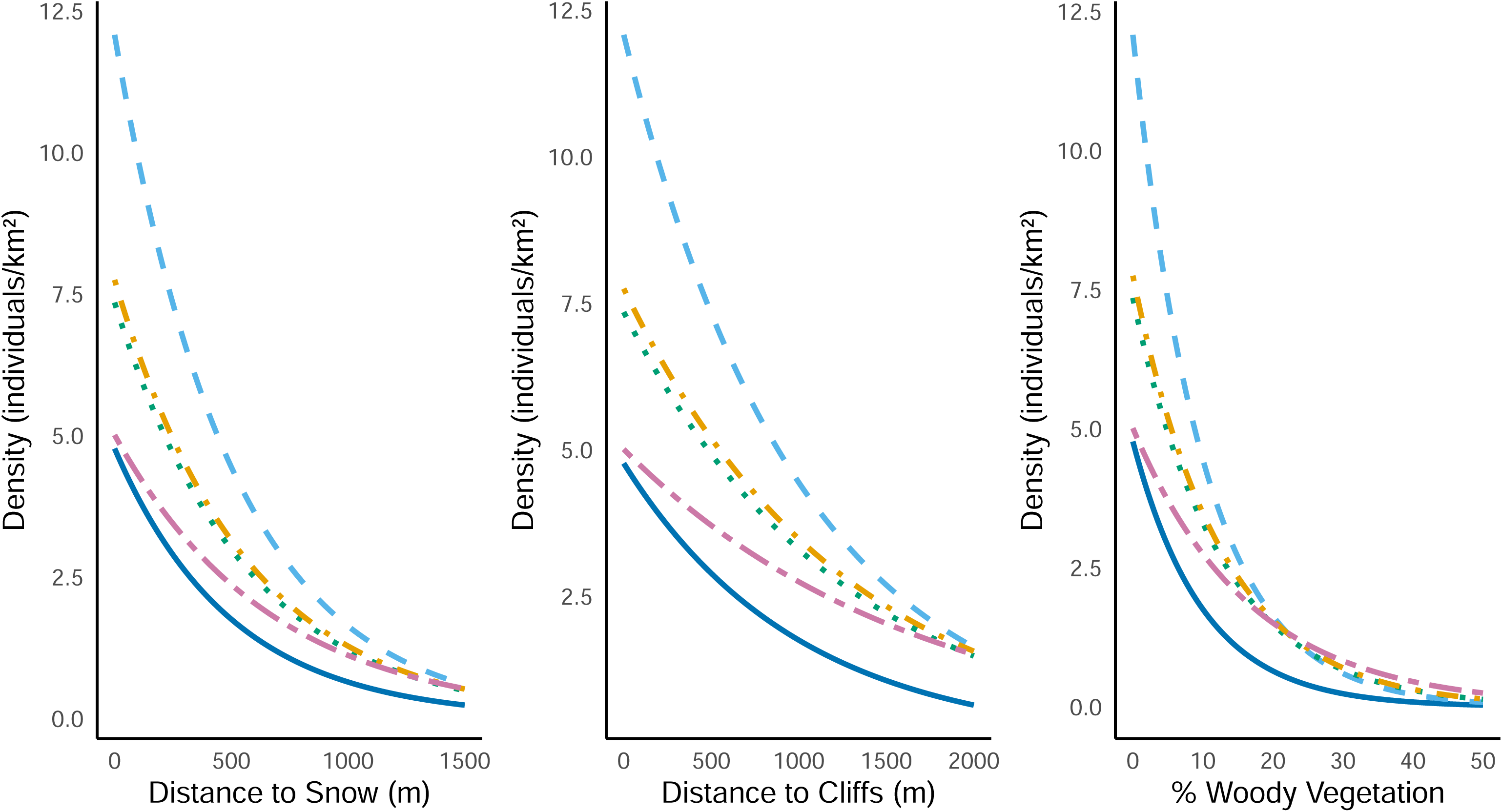
Predicted density of Sierra Nevada Rosy-Finch (*Leucosticte tephrocotis dawsoni*) as a function of three habitat covariates, 2018–2022. Response curves show model-predicted density (individuals km⁻²) from top-ranked hierarchical distance sampling models (ΔAIC ≤ 2) as functions of (a) distance to persistent snow patches (m), (b) distance to cliff faces (m), and (c) woody vegetation cover (%). All other covariates are held at their mean values. Line colors and styles distinguish years. Density declined consistently with increasing distance from snow and cliffs, and with increasing woody vegetation cover, across all five years. See Table 3 for model structures.

**Figure 3.**
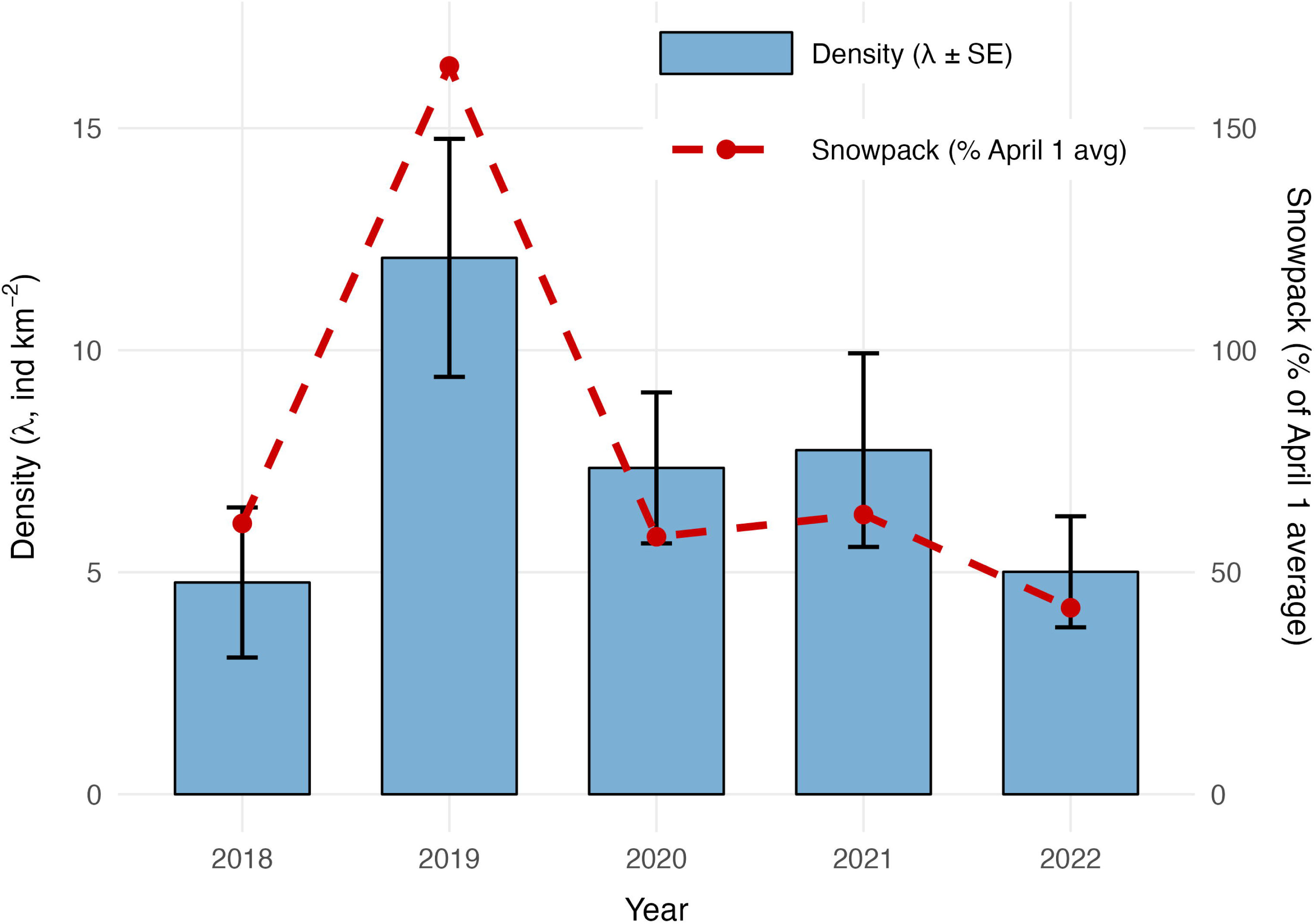
Predicted density of Sierra Nevada Rosy-Finch (*Leucosticte tephrocotis dawsoni*) as a function of three habitat covariates, 2018–2022. Response curves show model-predicted density (individuals km⁻²) from top-ranked hierarchical distance sampling models (ΔAIC ≤ 2) as functions of (a) distance to persistent snow patches (m), (b) distance to cliff faces (m), and (c) woody vegetation cover (%). All other covariates are held at their mean values. Line colors and styles distinguish years. Density declined consistently with increasing distance from snow and cliffs, and with increasing woody vegetation cover, across all five years. See Table 3 for model structures.

**TABLE 3.** Top-ranked hierarchical distance-sampling models (ΔAIC ≤ 2) for Sierra Nevada Rosy-Finch by year (2018–2022).

| Year | Covariates | K | AIC | $\Delta\text{AIC}$ | AICwt |
| --- | --- | --- | --- | --- | --- |
| 2018 | 1 ~ woody + snow + cliffs ($\lambda$) | 5 | 219.2 | 0.00 | 0.37 |
| 2018 | slope ($\sigma$) ~ woody + snow:cliffs ($\lambda$) | 6 | 219.5 | 0.31 | 0.32 |
| 2018 | time<sup>2</sup> ($\sigma$) ~ woody + snow:cliffs ($\lambda$) | 6 | 219.6 | 0.36 | 0.31 |
| 2019 | snow ($\sigma$) ~ woody + snow:cliffs + water ($\lambda$) | 7 | 479.9 | 0.00 | 1.00 |
| 2020 | slope ($\sigma$) ~ woody + snow + cliffs ($\lambda$) | 5 | 292.3 | 0.00 | 0.73 |
| 2020 | slope ($\sigma$) ~ woody + snow:cliffs ($\lambda$) | 6 | 294.3 | 1.94 | 0.27 |
| 2021 | 1 ~ woody + snow:cliffs ($\lambda$) | 5 | 375.8 | 0.00 | 0.47 |
| 2021 | time<sup>2</sup> ($\sigma$) ~ woody + snow:cliffs ($\lambda$) | 6 | 376.5 | 0.77 | 0.32 |
| 2021 | slope ($\sigma$) ~ woody + snow:cliffs ($\lambda$) | 6 | 377.4 | 1.59 | 0.21 |
| 2022 | slope ($\sigma$) ~ woody + snow + cliffs ($\lambda$) | 5 | 396.6 | 0.00 | 1.00 |

*Note:* Top-ranked models ( $\Delta\text{AIC} \leq 2$ ) are shown for each year. Covariates: Snow, distance to persistent snow patches; Cliffs, distance to cliff faces; Woody, woody vegetation cover; S×C, snow × cliff interaction; Day, Julian day; Slope, terrain slope; Time, time of day. Abbreviations: K, number of parameters; AIC, Akaike information criterion; $\Delta\text{AIC}$ , difference from top model; AICwt, AIC weight; $\lambda$ , abundance submodel; $\sigma$ , detection submodel.

Across all five years, top-ranked HDS models (ΔAIC ≤ 2) consistently included both distance to cliffs and distance to snow, with consistent effect directions (higher densities nearer cliffs and snow) and a negative effect of woody cover (Table 3; Fig. 2). Both cliff distance and snow distance appeared in the top-ranked model in every year, and both predictors remained within 4 AIC units of the top model across all candidate parameterizations in each year. Abundance declined sharply when woody cover exceeded ∼10%, with model-predicted densities reduced by more than 50% relative to open sites, and near-zero densities beyond ∼20–25% cover (Fig. 2), a threshold pattern consistent across all five years.

Annual detection records (Table S1) revealed consistent spatial variation in the proportion of surveyed blocks with detections. On average, 60.0% of blocks (range: 50.0–68.4%) contained detections each year, while a subset of blocks exhibited persistent non-detections (Table S1). In contrast, detection-corrected density estimates over the same period varied nearly threefold (4.77–12.08 individuals/km²; Table 2), a pattern that tracks interannual snowpack variation closely (Fig. 3) and illustrates why abundance provides resolution that block-level occupancy alone cannot.

## Discussion

Detection-corrected abundance patterns revealed sensitivity to both interannual snowpack variation and directional woody encroachment. Interpreted through a three-timescale co-limitation framework, in which breeding habitat is jointly constrained by static features (cliffs), dynamic annual drivers (snowpack), and longer-term directional change (woody encroachment), these results indicate that SN Rosy-Finch abundance is strongly associated with climate-related habitat variation within alpine systems. Our findings show that the abundance of the high-elevation specialist SN Rosy-Finch is governed by fine-scale spatial and temporal variation in climate-sensitive alpine resources, including stable abiotic features like cliffs and dynamic biotic drivers like snowpack. This resource dependence highlights the climate sensitivity of alpine vertebrates to habitat change driven by ongoing warming. While these associations indicate sensitivity to snowpack variability and woody encroachment, they do not preclude adaptive capacity. Sierra Nevada Rosy-Finches exhibit documented dietary flexibility, shifting among wind-deposited insects on snow, emergent aquatic insects, and alpine vegetation as resources become available (Epanchin et al. 2010; Clapp et al. 2025). Such trophic generalism may buffer short-term interannual variability in snowpack by allowing individuals to exploit alternative foraging substrates when persistent snow is limited. Additionally, fine-scale topographic heterogeneity across the Sierra Nevada creates localized zones of prolonged snow persistence and microclimatic buffering, potentially functioning as climate refugia (Morelli et al. 2016; Dobrowski 2011). Whether such refugia meaningfully sustain breeding populations under sustained warming remains an open question that fine-scale demographic and snow-persistence mapping could address. Recent genomic and morphometric analyses further suggest that alpine rosy-finch populations may exhibit signatures of local adaptation to elevational extremes (Robertson et al. 2026), indicating potential evolutionary capacity that warrants further investigation. These alternative mechanisms do not negate the strong, repeatable correlation between abundance and snowpack observed across five years and three mountain ranges, but instead highlight that the proximate pathways linking snowpack to population density remain an important area for future demographic investigation.

We observed heightened SN Rosy-Finch abundance during high-snow years (e.g., 2019; 162% of average snowpack) and marked declines during low-snow years (e.g., 2021–2022), indicating a strong correlation between population density and annual variation in snow availability (Fig. 3). The same-year correlation between snowpack and density warrants mechanistic consideration. Several non-exclusive explanations are plausible. During high-snow years, persistent snowfields may concentrate wind-deposited and emergent invertebrates along melt margins, increasing foraging opportunities and producing genuine increases in local density (Epanchin et al. 2010; Hodkinson et al. 2004). Alternatively, birds foraging on open snow may be more visually conspicuous than individuals using vegetated alpine substrates. However, SN Rosy-Finches are highly detectable during the breeding season. They are vocal, behaviorally bold, occur in open terrain at high elevations with low avian species richness, and are readily distinguished by plumage and call. Our MRDS analysis further supported high detectability within the effective radius (p(0) = 0.98), and slope was included as a detection covariate in the HDS models to account for terrain-related visibility differences. Percent snow cover was not explicitly included in the detection submodel. Ongoing spatial modeling efforts integrating remotely sensed snow metrics such as SWE and NDSI with detection-corrected density estimates will allow explicit evaluation of snow persistence across unsampled alpine habitat and strengthen inference about how climate-driven snow dynamics structure population distribution (e.g., Goljani Amirkhiz et al. 2025). Taken together, the strength and repeatability of the snow–density relationship across five years and three mountain ranges support an ecological association between snowpack and abundance, even as the proximate demographic pathways remain to be fully resolved.

Our findings align closely with early alpine surveys relating annual variation in the subspecies to snowpack variation (Grinnell & Storer 1924; Twining 1940). This century-long consistency between historical and contemporary patterns underscores the predictive value of snowpack as a leading indicator of alpine bird population sensitivity. However, continued evaluation of demographic rates, including overwinter survival and reproductive success will be necessary to determine the extent to which snowpack variability translates into long-term population change. Placed within this framework, static cliffs, annually variable snow, and directional woody encroachment, our five-year results generalize beyond a single species to other resource-limited, high-elevation taxa. Consistent with this view, the year with the greatest overdispersion in our models (2019; ĉ = 1.94) likely captured additional ecological heterogeneity associated with an extreme snow year, when unusually persistent snowpack may have altered movement patterns and detectability at fine scales (Table S2).

The spatial and temporal consistency of our results supports a framework in which SN Rosy-Finch populations are co-limited by fixed (cliffs) and annually variable (snow) resources, while facing increasing constraints from directional habitat change, particularly woody encroachment. This pattern mirrors trends observed in other alpine specialists (Beever et al. 2013; Resano-Mayor et al. 2019). Although occupancy remained relatively stable across years (range: 50.0–68.4% of blocks with detections; Table S1), variation in detection-corrected abundance tracked snowpack fluctuations nearly threefold across the same period (4.77–12.08 individuals/km²; Table 2; Fig. 3). This contrast is consistent with the expectation that abundance tracks habitat quality variation before occupancy declines (Sillett et al. 2012; Beever et al. 2013), and illustrates why abundance monitoring adds value in systems where populations may be declining in quality before declining in distribution. We note that this comparison is descriptive: a formal hierarchical occupancy model was not fitted, and we cannot rule out that an analogous detection-corrected occupancy analysis would reveal similar interannual variation. Nevertheless, the raw contrast between stable block-level occupancy and highly variable density estimates supports the value of abundance-based monitoring for early detection of climate-driven habitat change.

Collectively, the repeated selection of cliff and snow predictors across years, together with directionally consistent effects and year-dependent support for their interaction, indicates joint limitation by static (cliffs) and dynamic (snow) resources, while woody encroachment imposes a directional constraint. These findings highlight the value of integrating abundance monitoring into long-term alpine survey frameworks and contribute to a broader understanding of how alpine species respond to layered environmental pressures.

### Key habitat drivers and habitat associations

Persistent snow patches and rocky cliffs consistently emerged as the dominant drivers of SN Rosy-Finch abundance across years and regions, shaping the species’ fine-scale breeding distribution in California’s alpine ecosystems. Critically, the dominant drivers operate on distinct timescales: static (cliffs), annual (snow), and directional (woody encroachment). This distinction clarifies when conservation can act (limiting woody cover) versus when it must primarily monitor (snowpack). Our findings align with historical observations emphasizing the SN Rosy-Finch’s co-dependence on snowfields and rugged cliffs, features that often co-occur in high-elevation alpine landscapes and shape the species’ distribution (Twining 1940; Goljani Amirkhiz et al. 2025). Comparable cliff–snow associations have been reported for other rosy finch subspecies, including the Brown-capped Rosy-Finch (Johnson 1965), and are broadly documented in alpine bird communities worldwide (Scridel et al. 2018). Declining snowpack has been repeatedly linked to range contractions and reduced abundance (Beever et al. 2013; Goljani Amirkhiz et al. 2025), while woody vegetation encroachment alters alpine grassland habitats and may reduce habitat suitability for open-habitat specialists (Lubetkin et al. 2017; Scridel et al. 2018). Together, these results illustrate the three-timescale co-limitation framework: breeding habitat is constrained by static features (cliffs), dynamic annual drivers (snow), and long-term directional change (woody encroachment). This framework can be applied to other resource-limited systems to anticipate climate vulnerability.

Persistent snowfields and their mosaic boundaries with exposed alpine vegetation consistently supported SN Rosy-Finch detections, underscoring the importance of snowpack for reliable prey access during breeding. These zones concentrate emergent and wind-blown invertebrates along melting margins, serving as high-quality foraging habitat (Hodkinson et al. 2004; Epanchin et al. 2010; Brambilla et al. 2019; Resano-Mayor et al. 2019). Such mosaic snow–vegetation habitats facilitate elevated insect activity during key reproductive windows, as documented in snowfield-associated alpine passerines including the White-winged Snowfinch (*Montifringilla nivalis*) in the European Alps (Brambilla et al. 2019) and other high-elevation specialists across mountain systems (Scridel et al. 2018; Resano-Mayor et al. 2019). In contrast, cliffs offer stable nesting structures through time, often located on northeast-facing slopes where shading prolongs snow persistence. These sites reduce predation risk and buffer against thermal extremes during the breeding season (Pepin et al. 2015; Brown et al. 2018; Bernier et al. 2023). This dual reliance on dynamic and static features reflects adaptations to patchy, resource-limited alpine environments and mirrors habitat dependencies observed across other high-elevation specialists, including the White-winged Snowfinch (*Montifringilla nivalis*) in the European Alps (Brambilla et al. 2017; Scridel et al. 2018; Resano-Mayor et al. 2019). Our findings also support evidence that alpine birds shift spatially within breeding seasons in response to fine-scale snow and habitat dynamics, highlighting the value of temporally and spatially explicit monitoring (Resano-Mayor et al. 2019; Ceresa et al. 2020).

Strong negative associations between SN Rosy-Finch density and woody vegetation highlight the species’ sensitivity to directional shrub encroachment in alpine zones (Lubetkin et al. 2017; Scridel et al. 2018). Abundance declined sharply when woody cover exceeded ∼10%, with near-zero densities beyond ∼20–25% cover (Fig. 2), a threshold pattern consistent across all five years and all three mountain ranges sampled. This pattern is consistent with avoidance of structurally complex alpine patches, and we acknowledge that our observational design cannot distinguish between active displacement from previously occupied habitat and long-term disuse of areas where encroachment preceded our sampling window. Either mechanism would reduce the effective area of high-quality breeding habitat as woody cover increases. The apparent threshold indicates a functional reduction in breeding suitability as alpine vegetation structure becomes more complex. These findings emphasize a high degree of habitat specialization for open, low-structure landscapes. Similar threshold responses have been observed in other alpine specialists, such as the Black Rosy-Finch, which preferentially occupies snow-adjacent, rocky areas with minimal vegetation (Brown et al. 2018). These patterns are consistent with climate-driven treeline expansion, which increasingly compresses suitable habitat for alpine fauna (Körner 2003; Dirnböck et al. 2003; Pepin et al. 2015; Rumpf et al. 2019). Recent work in the European Alps confirms that shrub expansion is a key driver of alpine bird community turnover and replacement of cold-adapted species by generalists, even within protected areas (Alba et al. 2022; Alba & Chamberlain, 2025). Unlike snowpack variation, which fluctuates annually, woody encroachment represents a cumulative directional habitat change.

Taken together, our findings reveal that SN Rosy-Finch breeding habitat is shaped by a triad of interacting constraints operating across distinct temporal scales: cliffs provide stable nesting substrates, snowpack governs annual foraging opportunities, and woody encroachment exerts a slow-moving but directional threat to open alpine landscapes. This layered resource dependency underscores the species’ compounded vulnerability to climate change, with acute (interannual) fluctuations in snowpack intersecting with chronic (decadal) vegetation encroachment. These dual timescales of constraint reinforce the need for conservation strategies that address both dynamic and directional drivers of habitat loss. More broadly, this constitutes a general framework for diagnosing climate vulnerability in alpine specialists and other taxa confined to resource mosaics governed by multiple timescales.

### Regional variation in habitat use and comparative ecology

Comparisons with closely related taxa, the Brown-capped Rosy-Finch (*Leucosticte australis*) and Black Rosy-Finch (*Leucosticte atrata*), emphasize how regional climate differences shape alpine habitat specialization. In contrast to the Rocky Mountains, where higher and more consistent summer precipitation sustains stable foraging opportunities (Clow 2010; McGuire et al. 2012; Kittel et al. 2015; Bueno de Mesquita et al. 2018), California’s drier Mediterranean alpine climate results in patchier, less predictable snowfields (Siirila-Woodburn et al. 2021). This climate-driven variability forces SN Rosy-Finches to range more extensively from nesting cliffs and snow patches to access dispersed resources, likely increasing energetic costs and reducing breeding success relative to their Rocky Mountain counterparts (Brown et al. 2018; Bernier et al. 2023). This behavioral pressure could plausibly drive selection on morphometric traits associated with longer-range foraging, such as wing shape or body mass, though future comparative studies are needed to evaluate this possibility. Bernier et al. (2023) explicitly documented that Brown-capped Rosy-Finches benefited from stable snowfields and higher insect productivity linked to more consistent summer rainfall, contrasting sharply with the ephemeral resource conditions typically found in California’s alpine regions. Stable snow and predictable melt timing regulate alpine invertebrate pulses that underpin passerine foraging during breeding (Morton 1978; Antor 1995; Epanchin et al. 2010; Brambilla et al. 2019). Comparable fine-scale preferences are reported for alpine northern wheatear (*Oenanthe oenanthe*) in the Alps, which selects open, short-sward, snow-proximate microhabitats and avoids woody cover, underscoring the importance of accessible foraging substrates under climate and land-use change (Müller et al. 2023).

Intra-regional comparisons further reveal that SN Rosy-Finch habitat use varies with aridity, reinforcing patterns of increased energetic cost and vulnerability in drier landscapes. In the White and Sweetwater Mountains, more arid than the Sierra Nevada, finches consistently foraged farther from key habitat features (median distances: 400 m from cliffs vs. 150 m; 350 m from snow vs. 100 m; Fig. S4). Persistent non-detections in our dataset were concentrated disproportionately in the White Mountains and Sweetwater Mountains relative to the Sierra Nevada (Table S1), consistent with the lower predicted habitat suitability and more contracted range documented in those ranges by Goljani Amirkhiz et al. (2025). Figure S4 suggests that habitat associations with cliffs and snow may differ in magnitude across mountain ranges, though our sampling within the White and Sweetwater Mountains was insufficient to formally test this. Expanded spatial coverage combined with range-wide density modeling will allow evaluation of whether these habitat relationships shift along aridity gradients or remain consistent across the subspecies’ distribution. These longer travel distances may impose higher metabolic costs and reduce habitat efficiency, especially in fragmented terrain. When cliffs and transient snowfields do not occur within energetically efficient commuting distances, longer travel times can reduce chick provisioning and lower breeding success (Fig. S4; Orians & Pearson 1979; Pyke 1984). Historical observations compiled by Goljani Amirkhiz et al. (2025) corroborate this, revealing substantial range contractions over the last 80 years that appear linked to warming-induced habitat loss. Together, these results emphasize that climate impacts on SN Rosy-Finch are not uniform across its range, but intensify in the most arid and topographically fragmented portions.

### Methodological considerations

Our approach combined Mark-Recapture Distance Sampling (MRDS) to estimate observer detection probabilities with Hierarchical Distance Sampling (HDS) to estimate abundance. HDS models produced the site-specific, detection-corrected density estimates (Table 2), while MRDS validated the assumption of high detection and minimal observer bias (Table 1, Fig. S2). These models explicitly accounted for detection biases and spatial heterogeneity, common challenges in bird research (Buckland et al. 2001; Sillett et al. 2012; Kéry & Royle 2015), and built on established practices for estimating habitat-specific abundance in challenging environments. Our staggered-observer implementation of MRDS adapted field protocols to the rugged terrain of the Sierra and still yielded consistent detection probabilities (0.88 and 0.86), reinforcing the species’ high detectability. GLMM analyses complemented this approach by elucidating spatial random effects (block-level clustering) and further informing variable selection in subsequent HDS models. However, slight overdispersion associated with some models (ĉ values up to 1.94; Table S2) was within acceptable ecological modeling ranges (ĉ < 2) and points to additional unmodeled heterogeneity, potentially arising from unmeasured habitat variables or behavioral factors. Future studies could incorporate additional habitat covariates (e.g., microtopographic variability, prey abundance), such as snowmelt timing derived from satellite imagery or fine-scale terrain complexity metrics like Vector Ruggedness Measure (VRM), to further refine model accuracy. As with all survey-based models, inference is bounded by the spatial and temporal extent of sampling. Future efforts to integrate abundance models with spatial prediction frameworks (e.g., occupancy-abundance SDMs) could improve extrapolation capacity and inform range-wide conservation strategies.

The HDS framework enabled detection of interannual climate sensitivity, with nearly threefold variation in density tracking snowpack, that block-level occupancy did not reveal (range: 50.0–68.4% across years; Table S1), consistent with the expectation that abundance declines precede occupancy loss in contracting populations (Beever et al. 2013; Sillett et al. 2012). This distinction has direct methodological implications, as monitoring programs relying solely on presence/absence data for this subspecies would be less likely to detect the signal documented here.

### Establishing baselines for conservation and monitoring

High-elevation alpine ecosystems are underrepresented in avian surveys, resulting in critical knowledge gaps (Siegel et al. 2014; Tingley et al. 2009). While Tingley et al. (2009) documented climate-driven elevational shifts across Sierra Nevada birds, logistical constraints limited sampling in the highest alpine zones where SN Rosy-Finches obligately breed, and contemporary alpine bird studies, both in North America and Europe (Brambilla et al. 2019; Resano-Mayor et al. 2019), have typically relied on occupancy rather than detection-corrected density estimates.

Our study addresses these gaps by providing a range-wide, detection-corrected, abundance-informed baseline for SN Rosy-Finch densities and habitat relationships, combining fine-scale field data with rigorous detection-corrected hierarchical modeling frameworks. This approach not only offers a replicable template for long-term ecological monitoring, but also solidifies essential context for future landscape-scale assessments using species distribution models (SDMs) and GIS-based habitat suitability mapping to inform refined population estimates and conservation prioritization. In doing so, it establishes the empirical foundation for forecasting persistence via spatially explicit population and distribution modeling, an immediate next step for management-relevant scenario testing.

Taken together, our findings establish a detection-corrected, site-specific ecological baseline with methodological and conceptual relevance beyond a single-species case study. We distinguish carefully between pattern and mechanism. Our observational framework identifies the habitat correlates of abundance, including cliffs, snowpack, and woody cover, but does not resolve the physiological, behavioral, or evolutionary pathways by which these features shape population dynamics. This work provides rigorously derived, detection-corrected density benchmarks and clearly defined habitat associations that represent the sensitivity and exposure components of a broader climate vulnerability assessment for this subspecies, as well as a transferable analytical framework for diagnosing similar resource constraints in other alpine taxa. It also lays the groundwork for future comparative studies across alpine bird communities and highlights the value of high-resolution, detection-corrected monitoring for forecasting species vulnerability under ongoing climate change.

Our findings about SN Rosy-Finch have broader ecological relevance and implications for conservation strategies in alpine systems worldwide. Similar to high-elevation specialists like the American pika (Beever et al. 2013), SN Rosy-Finches illustrate how abundance-based approaches expose vulnerability to multiple, interacting habitat constraints often masked by simpler presence/absence metrics. These parallels reinforce the value of abundance-based approaches for detecting climate-sensitive responses in alpine specialists and support their integration into long-term monitoring frameworks.

### Climate vulnerability, adaptive potential, and conservation implications

The strong link between SN Rosy-Finch densities and snowpack variability underscores this species’ pronounced vulnerability to climate-driven habitat changes. Population declines following low-snow years, particularly in 2021–2022, may reflect both immediate demographic impacts and cumulative lag effects (e.g., overwinter mortality, reduced reproductive success) from harsh conditions. While the correlation between annual snowpack and breeding-season density is strong (Fig. 3), we cannot fully disentangle whether this reflects within-season habitat quality (persistent snowfields providing foraging resources during surveys), behavioral aggregation (birds concentrating in high-quality habitat during high-snow years), or lagged demographic effects from prior-year reproductive success. Future studies should explicitly disentangle these drivers through demographic analyses (e.g., overwinter survival, fledgling success), which will be critical for forecasting population trends and informing responsive conservation strategies.

Given that SN Rosy-Finch already occupies the highest available elevations and faces limited potential for range shifts, conservation efforts should emphasize habitat protection and enhancement within existing alpine zones. Continued warming and associated elevational range contractions threaten restricted alpine habitats globally (Freeman et al. 2018). Brown et al. (2018) documented similar vulnerability for the Black Rosy-Finch, underscoring that upward shifts driven by climate warming pose severe constraints for alpine species. This is a critical concern for the SN Rosy-Finch. Genomic analyses reveal significant genetic differentiation between the Sierra Nevada population and other Gray-crowned Rosy-Finch subspecies, with evidence of local adaptation to the distinctive elevational and climatic conditions of the Sierra Nevada (Robertson et al. 2026). This elevates the conservation urgency of this distinct alpine lineage in the face of accelerating climate impacts.

### Future directions for alpine bird conservation

Long-term, multi-scale monitoring that integrates genetic data and remote sensing (e.g., LiDAR, satellite imagery) will be crucial to assess habitat connectivity, fragmentation risks, and species’ adaptive responses to environmental change. European studies on alpine specialists like the White-winged Snowfinch underscore the effectiveness of combining fine-scale ecological data with broad-scale remote sensing for understanding and mitigating climate impacts (Brambilla et al. 2017; Resano-Mayor et al. 2019). Such integrative methods can guide proactive conservation strategies tailored to alpine species facing accelerated habitat loss, and implementable by land managers and monitoring networks.

Ultimately, effective conservation of alpine birds requires preserving open, low-vegetation habitats characterized by persistent snow patches and stable nesting sites. Achieving this will necessitate coordinated, landscape-scale management interventions specifically designed to mitigate woody vegetation encroachment, address compounding vulnerabilities (e.g., annual snow reduction, long-term shrub invasion), and reduce habitat fragmentation. The ecological baselines established by our study, namely, spatially explicit abundance estimates and clearly defined habitat associations provide a critical foundation for these proactive strategies. By establishing abundance baselines linked to climate-sensitive resources, this approach provides a template for climate vulnerability assessments across high-elevation taxa worldwide.

## Authorship contributions

Timothy Brown contributed to conceptualization, study design, fieldwork coordination, data collection, visualization, formal analysis, writing the original draft, and reviewing and editing. Erika Zavaleta contributed to conceptualization, study design, methodology, funding acquisition, project administration, and reviewing and editing. Reza Goljani Amirkhiz provided methodological guidance, model development, analytical strategy, and reviewing and editing. Kristen Ruegg and Mevin Hooten contributed to funding acquisition and reviewing and editing. Kathryn Bernier provided conceptual input and scientific review. T.B., E.Z., R.G.A., K.R., M.H., and K.B. discussed the results and contributed to the final manuscript. Sarah Albright, Alena Arnold, Emma Brown, Carl Brown, Vincent Chevreuil, Ryan Cheung, Daisy Cortes, Jessica Gallardo, Karim Hanna, Raquel Lozano, Juan Rebellon, Leticia Santillana, Kevin Silberberg, and Juyung Yoo provided field support and data collection assistance.

## Acknowledgments

We thank the Conservation Science & Stewardship Lab at the University of California, Santa Cruz, for their invaluable support and feedback throughout this project. Special thanks to Sushmita Poudel, Abe Borker, and Kelly Zilliacus for their insights and encouragement. We are especially grateful to Bruce Lyon and Pete Raimondi (UCSC) for expert guidance on ecological interpretation, statistical analyses, and methodological rigor. We deeply appreciate Yosemite and Sequoia–Kings Canyon National Parks and the National Park Service for continued support and permitting assistance, and the U.S. Forest Service for facilitating access to field sites within Inyo, Sierra, Stanislaus, and Humboldt–Toiyabe National Forests. This project was supported by the National Science Foundation (Grant Nos. 2222524, 2222525, 2222526), the Mildred E. Mathias Graduate Student Research Grant (University of California Natural Reserve System), the Sequoia Parks Conservancy (the official nonprofit partner to Sequoia and Kings Canyon National Parks), and the Bloom-Hays Ecological Research Grant (Sea and Sage Audubon Society).

