## Supporting Information for "Co-limitation by stable, dynamic and directional habitat features shapes climate vulnerability in an alpine specialist"

Ecological Applications

**TABLE S1** Annual survey effort, detections, and block-level occupancy for Sierra Nevada Rosy-Finch, 2018–2022.

| Year | Blocks surveyed | Survey stations | Detections | Blocks with detections | Occupancy (%) |
|---|---|---|---|---|---|
| 2018 | 16 | 128 | 72 | 8 | 50.0 |
| 2019 | 19 | 152 | 119 | 13 | 68.4 |
| 2020 | 19 | 152 | 81 | 10 | 52.6 |
| 2021 | 21 | 168 | 86 | 14 | 66.7 |
| 2022 | 21 | 168 | 102 | 13 | 61.9 |

*Note:* Blocks represent 4 × 4 km sampling units. Occupancy (%) is the percentage of blocks with ≥1 detection. Abbreviation: GCRF, Gray-crowned Rosy-Finch.

**TABLE S2** Goodness-of-fit metrics for hierarchical distance sampling models, 2018–2022.

| Year | SSE | FT residuals | c-hat |
|---|---|---|---|
| 2018 | 49.5 | 44.4 | 1.31 |
| 2018 | 47.7 | 43.8 | 1.60 |
| 2018 | 49.1 | 43.6 | 1.32 |
| 2019 | 123.0 | 108.0 | 1.79 |
| 2020 | 63.5 | 62.7 | 1.01 |
| 2020 | 63.5 | 62.7 | 1.04 |
| 2021 | 72.0 | 77.0 | 1.45 |
| 2021 | 72.0 | 76.7 | 1.35 |
| 2021 | 71.9 | 77.5 | 1.41 |
| 2022 | 93.1 | 86.3 | 1.51 |

*Note:* c-hat values near 1 indicate adequate model fit; values >1 suggest overdispersion. Abbreviations: SSE, sum of squared errors; FT, Freeman–Tukey residuals; c-hat, overdispersion estimate.

**TABLE S3** Coefficient estimates from hierarchical distance sampling models of Sierra Nevada Rosy-Finch density and detection probability, 2018–2022.

| Year | Component | Parameter | Estimate | SE | Z | P | Sig. | CI low | CI high |
|---|---|---|---|---|---|---|---|---|---|
| 2018 | Abundance | Intercept | -5.34 | 2.21 | -2.41 | 0.0159 | * | -9.68 | -1.00 |
| 2018 | Abundance | **Woody** | -4.81 | 2.67 | -1.80 | 0.0717 | | -10.05 | 0.43 |
| 2018 | Abundance | **Dist_Snow** | -4.08 | 1.22 | -3.36 | 0.0008 | *** | -6.46 | -1.70 |
| 2018 | Abundance | **Dist_Cliffs** | 1.41 | 0.44 | 3.23 | 0.0013 | ** | 0.55 | 2.27 |
| 2018 | Detection | Intercept | 4.30 | 0.14 | 29.70 | 0.0000 | *** | 4.02 | 4.58 |
| 2018 | Dispersion | Dispersion | -2.17 | 0.42 | -5.13 | 0.0000 | *** | -3.00 | -1.34 |
| 2019 | Abundance | Intercept | -10.20 | 6.00 | -1.70 | 0.0892 | | -21.96 | 1.56 |
| 2019 | Abundance | **Woody** | -1.33 | 0.45 | -2.97 | 0.0030 | ** | -2.21 | -0.45 |
| 2019 | Abundance | **Dist_Snow** | -10.01 | 6.74 | -1.48 | 0.1377 | | -23.22 | 3.20 |
| 2019 | Abundance | **Dist_Cliffs** | -6.92 | 7.57 | -0.92 | 0.3604 | | -21.75 | 7.91 |
| 2019 | Abundance | Dist_Water | -2.63 | 1.02 | -2.58 | 0.0099 | ** | -4.63 | -0.63 |
| 2019 | Abundance | Snow:Cliffs | -8.02 | 8.65 | -0.93 | 0.3538 | | -24.97 | 8.93 |
| 2019 | Detection | Intercept | 4.18 | 0.10 | 41.00 | 0.0000 | *** | 3.98 | 4.38 |
| 2019 | Dispersion | Dispersion | -0.15 | 0.43 | -0.36 | 0.7200 | | -0.99 | 0.68 |
| 2020 | Abundance | Intercept | -5.97 | 1.69 | -3.53 | 0.0004 | *** | -9.28 | -2.66 |
| 2020 | Abundance | **Woody** | -4.29 | 2.08 | -2.06 | 0.0391 | * | -8.36 | -0.22 |
| 2020 | Abundance | **Dist_Snow** | -1.59 | 0.62 | -2.58 | 0.0098 | ** | -2.80 | -0.38 |
| 2020 | Abundance | **Dist_Cliffs** | -2.60 | 1.12 | -2.33 | 0.0197 | * | -4.79 | -0.41 |
| 2020 | Detection | Slope | -0.40 | 0.11 | -3.62 | 0.0003 | *** | -0.61 | -0.18 |
| 2020 | Dispersion | Dispersion | 0.76 | 0.84 | 0.91 | 0.3650 | | -0.88 | 2.40 |
| 2021 | Abundance | Intercept | -4.67 | 1.75 | -2.66 | 0.0078 | ** | -8.10 | -1.23 |
| 2021 | Abundance | **Woody** | -5.13 | 2.24 | -2.29 | 0.0220 | * | -9.51 | -0.74 |
| 2021 | Abundance | **Dist_Snow** | -2.06 | 0.81 | -2.56 | 0.0106 | * | -3.63 | -0.48 |
| 2021 | Abundance | **Dist_Cliffs** | -0.86 | 0.60 | -1.43 | 0.1534 | | -2.05 | 0.32 |
| 2021 | Abundance | Snow:Cliffs | -1.36 | 0.93 | -1.46 | 0.1447 | | -3.19 | 0.47 |
| 2021 | Detection | Intercept | 4.29 | 0.11 | 38.20 | 0.0000 | *** | 4.07 | 4.51 |
| 2021 | Dispersion | Dispersion | -0.52 | 0.48 | -1.07 | 0.2830 | | -1.46 | 0.43 |
| 2022 | Abundance | Intercept | -5.39 | 1.57 | -3.43 | 0.0006 | *** | -8.47 | -2.31 |
| 2022 | Abundance | **Woody** | -5.37 | 2.04 | -2.63 | 0.0086 | ** | -9.37 | -1.36 |
| 2022 | Abundance | **Dist_Snow** | -0.14 | 0.47 | -0.29 | 0.7699 | | -1.06 | 0.79 |
| 2022 | Abundance | **Dist_Cliffs** | -1.52 | 0.57 | -2.66 | 0.0077 | ** | -2.63 | -0.40 |
| 2022 | Detection | Slope | -0.38 | 0.20 | -1.93 | 0.0542 | | -0.77 | 0.01 |
| 2022 | Dispersion | Dispersion | -0.54 | 0.46 | -1.18 | 0.2400 | | -1.43 | 0.36 |

*Note:* Parameters in boldface indicate key habitat covariates (distance to cliffs, distance to snow, woody vegetation cover). All covariates were Z-standardized (mean = 0, SD = 1). Abbreviations: SE, standard error; CI, confidence interval; $\lambda$, abundance submodel; $\sigma$, detection submodel. Significance: * $P < 0.05$; ** $P < 0.01$; *** $P < 0.001$.

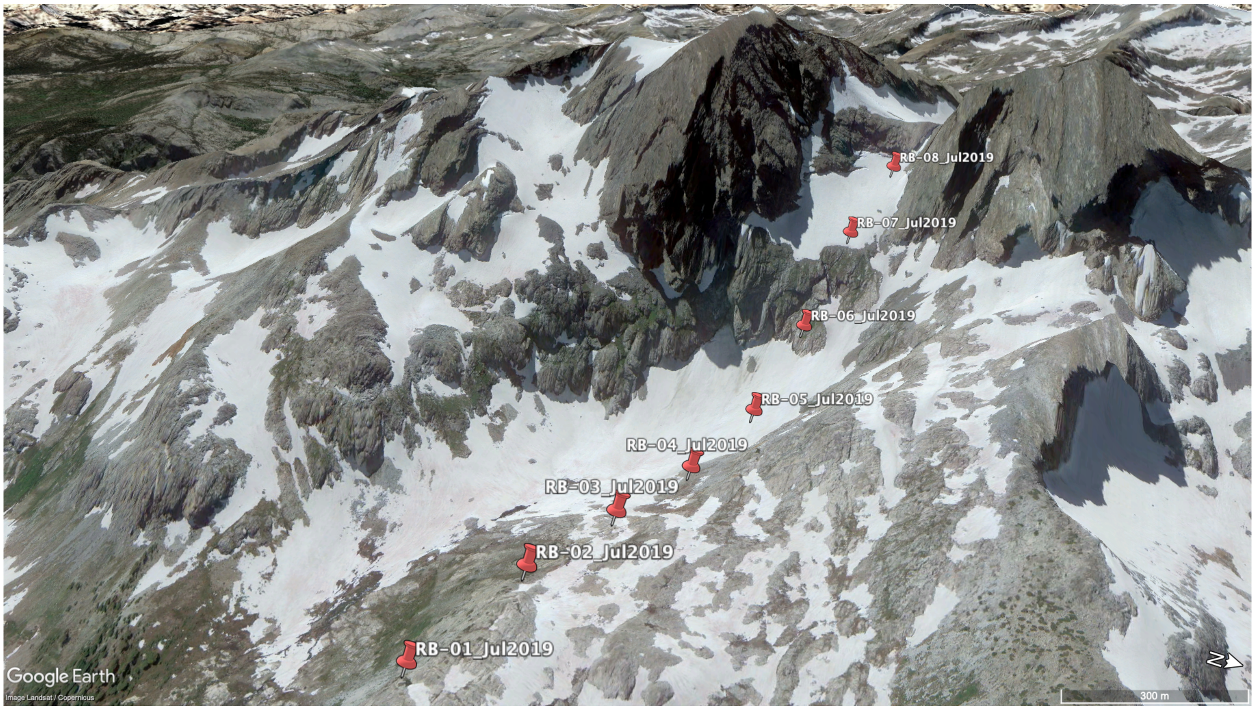


**FIGURE S1** Oblique aerial view of eight survey stations (RB-01 through RB-08, red markers) positioned every 250 m across alpine terrain between Mt. Ritter and Banner Peak in the Minarets Range, Sierra Nevada, surveyed July 2019. The landscape illustrates co-occurring habitat features critical to Sierra Nevada Rosy-Finch ecology: cliff faces (nesting substrate), persistent snowfields (foraging habitat), and patchy alpine vegetation. Scale bar: 300 m. Image: © Google Earth, Landsat/Copernicus.

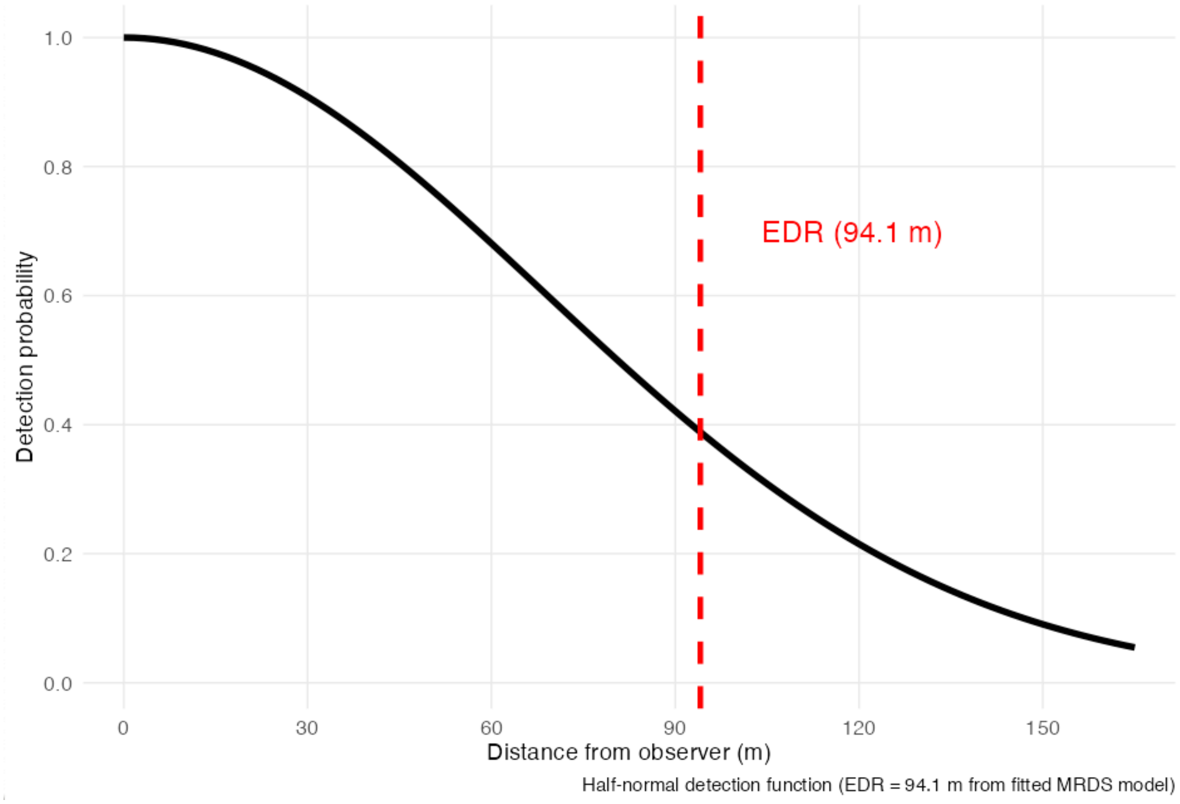


**FIGURE S2** Detection function derived from mark–recapture distance sampling (MRDS) analysis of Sierra Nevada Rosy-Finch detections, truncated at 165 m. The fitted half-normal detection function shows declining detection probability with increasing distance from the observer, consistent with expectations for point-transect surveys. The dashed red line indicates the effective detection radius (EDR = 94.1 m), the distance within which detection is most reliable. Observer-specific detection probability was $p_0 = 0.87$ and combined detection probability at distance zero was $p(0) = 0.98$ (95% CI: 0.96–0.99); parameter uncertainty was estimated using the model-based variance estimator. Abbreviations: EDR, effective detection radius; MRDS, mark–recapture distance sampling; $p_0$, single-observer detection probability; $p(0)$, combined detection probability at distance zero.

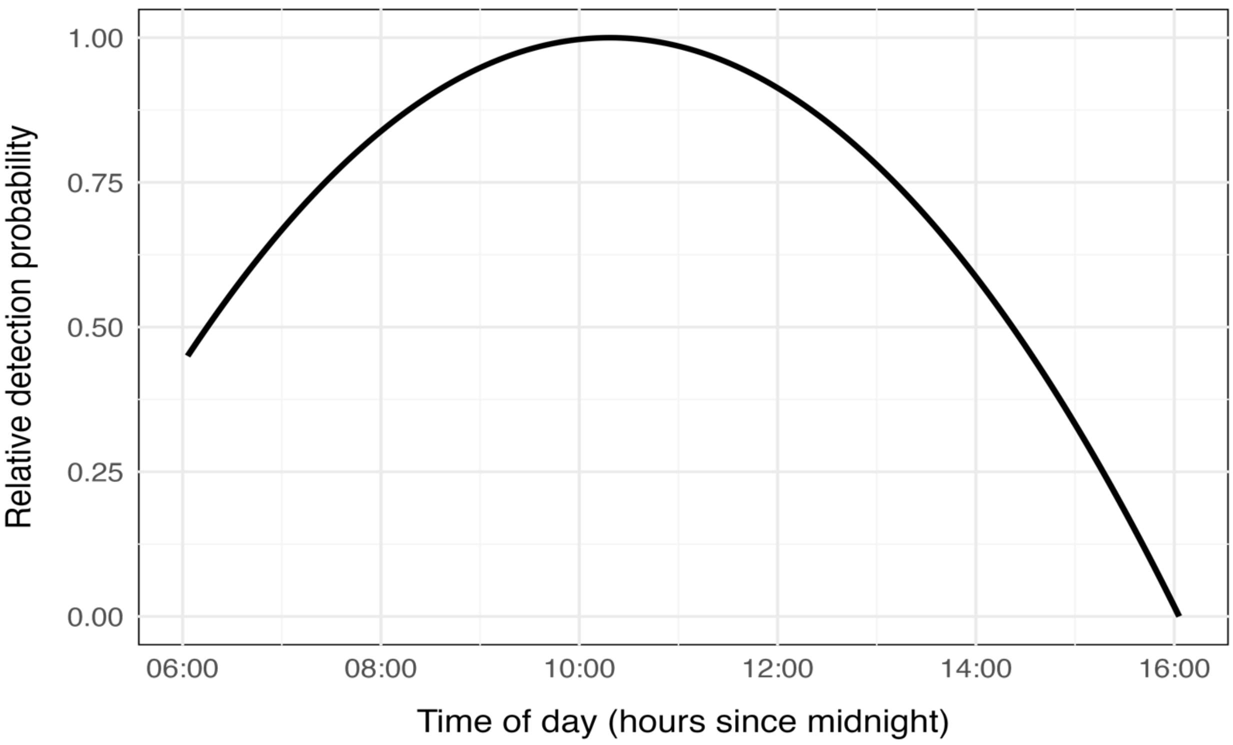


**FIGURE S3** Modeled detection probability of Sierra Nevada Rosy-Finch as a function of time of day, averaged across survey years. The quadratic detection curve peaks during mid-morning surveys (approximately 09:00–10:30) and declines through midday. Model predictions are restricted to the observed survey time range (06:00–13:00 h); axis labels extending beyond 13:00 are shown for visual reference only and do not represent model extrapolation. Surveys were completed by 13:00 h in all years.

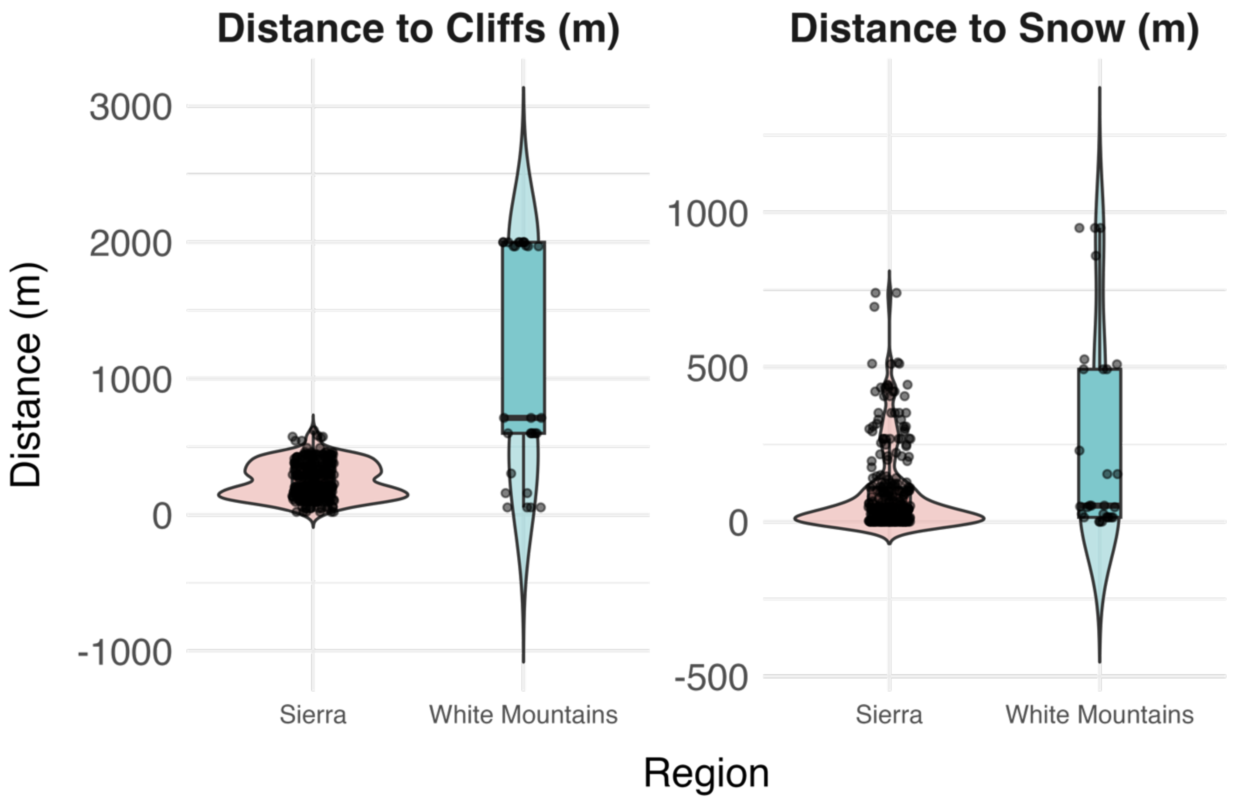


**FIGURE S4** Regional differences in habitat use by Sierra Nevada Rosy-Finch based on distances to cliffs and snowfields in the Sierra Nevada (blue) and White Mountains (red). Violin plots show the distribution of values, with boxplots indicating medians and interquartile ranges, and points representing individual observations. Birds in the Sierra Nevada were generally closer to cliffs and snow than those in the White Mountains, highlighting regional differences in alpine habitat use.
